# Evaluating analgesic efficacy and administration route following craniotomy in mice using the grimace scale

**DOI:** 10.1101/408443

**Authors:** Chulmin Cho, Vassilia Michalidis, Irene Lecker, Chereen Collymore, David Hanwell, Mary Loka, Matthew Danesh, Christine Pham, Paige Urban, Robert P. Bonin, Loren J. Martin

## Abstract

Most research laboratories abide by guidelines and mandates set by their research institution regarding the administration of analgesics to control pain during the postoperative period. Unfortunately, measuring pain originating from the head is difficult, making adequate decisions regarding pain control following stereotaxic surgery problematic. In addition, most postsurgical analgesia protocols require multiple injections over several days, which may cause stress and distress during a critical recovery period. Here we sought to 1) assess the degree of postoperative pain following craniotomy in mice, 2) compare the efficacy of three common rodent analgesics (carprofen, meloxicam and buprenorphine) for reducing this pain and 3) determine whether the route of administration (injected or self-administered through the water supply) influenced pain relief post-craniotomy. Using the mouse grimace scale (MGS), we found that injectable analgesics were significantly more effective at relieving post-craniotomy pain, however, both routes of administration decreased pain scores in the first 24 h post-surgery. Specifically, buprenorphine administered independently of administration route was the most effective at reducing MGS scores, however, female mice showed greater sensitivity to carprofen when administered through the water supply. Although it is necessary to provide laboratory animals with analgesics after an invasive procedure, there remains a gap in the literature regarding the degree of craniotomy-related pain in rodents and the efficacy of alternative routes of administration. Our study highlights the limitations of administering drugs through the water supply, even at doses that are considered to be higher than those currently recommended by most research institutions for treating pain of mild to moderate severity.

## Introduction

In neuroscience, there are a variety of approaches and techniques that require direct access to the rodent brain. Consequently, it has become increasingly common for many neuroscience labs to perform craniotomies, an invasive brain surgery in which the brain is accessed via removal of a section of the skull, so that studies involving intracranial injections, cannulations, electrical stimulation, or optical implants can be conducted ^1-3^. Following many of these surgeries, post-operative analgesics are often administered. However, the management of post-craniotomy pain has yet to be standardized and there are no current evidence-based recommendations for the alleviation of craniotomy pain in mice. Given that the insufficient management of acute pain following surgery may lead to depression 4,5 and anxiety – behaviors directly studied by many neuroscience labs – finely-tuned pain control following craniotomy should be prioritized and properly evaluated.

The establishment of a standardized analgesic regiment for post-craniotomy pain in animals has been hindered by the difficulty in assessing the degree of pain post-surgery and the efficacy of chosen analgesics. Traditional pain measurement in laboratory rodents has heavily relied on evoked pain behaviors, however, this becomes impractical for assessing pain originating from surgical operations involving the head and scalp (see 6 for a review). Non-selective proxy measures such as cardiovascular changes, food and water intake, locomotion, and nest construction have been used to measure ‘spontaneous’ pain following injury with varying degrees of success and large inter-subject variability^7-9^. In addition, learning paradigms such as conditioned place preference have also been used to evaluate postoperative pain and pain relief following surgery^10^, but these studies are often limited by one-trial learning making the assessment of pain over time difficult.

The recent implementation of the grimace scales has expanded our ability to assess pain in rodents^11,12^ and potentially addresses some of the issues associated with evoked or proxy measures of pain in laboratory animals ^6^. The grimace scale was originally developed for mice based on the facial action coding system for pain in infants and non-verbal humans ^13^. The mouse grimace scale (MGS), as it has come to be known, measures changes in facial musculature – orbital tightening, nose bulge, cheek bulge, ear change, and whisker change – following injury11. Each feature is given a score of 0 (not present), ^1^ (moderate) or 2 (severe), which can be scored with remarkably high reliability and accuracy. Although the grimace scale has been readily evaluated in mice ^11,14,15^ and rats ^12,16^, other grimace scales have been developed to capture ‘pain faces’ in horses ^17^, cats ^18^, rabbits ^19^, sheep ^20^ and ferrets ^21^, which have helped to evaluate postoperative pain across species.

The sensitivity of the MGS allows for the detection of postoperative pain upwards of 48 h following surgery – time points beyond this have not been assessed – and indicates that commonly used analgesics may not be efficacious at recommended doses ^14^. There is a strong positive correlation between MGS scores and pain-associated behaviors in mice ^22^, suggesting that facial expressions of pain could provide a rapid and reliable clinical pain assessment modality. Since the mouse grimace scale has been shown to be a reliable indicator of pain postsurgery ^14,15,22^, we used it as an assessment tool to characterize the extent and duration of postoperative pain in mice following craniotomy. We evaluated the efficacy of common rodent analgesics: the µ-opioid receptor partial agonist buprenorphine and nonsteroidal anti-inflammatory drugs (NSAID) carprofen and meloxicam. Further, since multiple injections of analgesics are often required for pain management in rodents 5, we assessed whether self-administration of these analgesics (i.e. through the water supply) offered comparable analgesia to the injectable formulations. We largely find that analgesics dministered through an injection route were more effective in mitigating spontaneous pain behavior when compared with free access through the water supply. Buprenorphine was the most effective for treating pain in both males and females, although minor sex differences emerged for carprofen.

## Results

A five-way ANOVA of all data (site x sex x administration route x drug x repeated measure) revealed a main effect for time course (F_5,142_=143.4, p<0.001), a main effect for administration route (F_1,142_=4.77, p=0.03) and a three-way interaction between sex, administration route and drug (F_5,142_=2.56, p=0.03). There was also a three-way interaction between administration route, drug and time course (F_25,142_=2.51, p<0.001). No other meaningful interactions reached significance (All F’s < 2.2, p>0.05). Since there was no main effect for site (F_1,142_=2.0, p=0.16) and analysis of baseline MGS scores did not reveal a difference between testing sites (*Martin Lab*, 0.07 × 0.01 vs. *Bonin Lab*, 0.07 × 0.01; mean × SEM, two-tailed *t-test*, *t*_188_=0. 99, *p*=0.3), data from both labs were combined. However, data were separated according to route of administration (injected vs water supply) and sex for analysis of the entire postoperative pain period (i.e. 72 h) based on the three-way interactions.

### Analysis of injected analgesics

Mean difference scores for each time point (MGS postoperative time point score – MGS baseline score) were calculated as previously done 14. The interaction between drug and time point was significant (two-way ANOVA, drug x repeated measures: F_25,88_=4.75, p<0.001) and posthoc testing (one-way ANOVA at each time point) showed that buprenorphine, carprofen (25 mg/kg) and meloxicam (5 mg/kg) reduced MGS scores compared with saline control across time points within the first 24 hrs following craniotomy (4 h, F_5,88_= 7.517, p<0.001; 6 h, F_5,88_=10.63, p<0.001; 8 h, F_5,88_=4.27, p=0.002; 24 h, F_5,88_=5.259, p<0.001). Carprofen (10 mg/kg; p=0.002) and meloxicam (2 mg/kg; p<0.001) had a slower onset, but by 6 h following craniotomy MGS scores were significantly reduced compared with control mice (**Figure 2A**). No drugs reduced MGS scores at 48 h (F_5,88_=0.93, p>0.05) or 72 h (F_5,88_=1.03, p>0.05) postsurgery even though scores in the saline group were significantly higher than 0 at 48 h postsurgery (t_15_=2.12, ***p=0.048)***. There were no overall sex differences for the injected analgesics (F_1,82_=0.072, p=.789), however, data were separated and plotted for male (**Figure 2B**) and female (**Figure 2C**) mice over the entire postoperative observation period.

**Figure 1.**
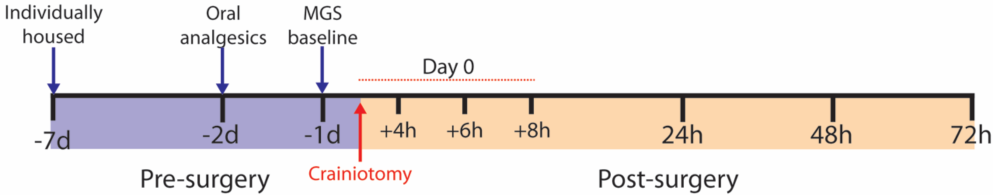
Experimental timeline outlining the pre- and postsurgical periods. For all mice, tap water was replaced with the MediDrop(r) solution 48 hrs prior to surgery. Analgesic drugs were then added to drinking water, as appropriate, beginning 48 hrs prior to surgery. MGS baseline scores were recorded 24 h before craniotomy and at the indicated time points following surgery.

**Figure 2.**
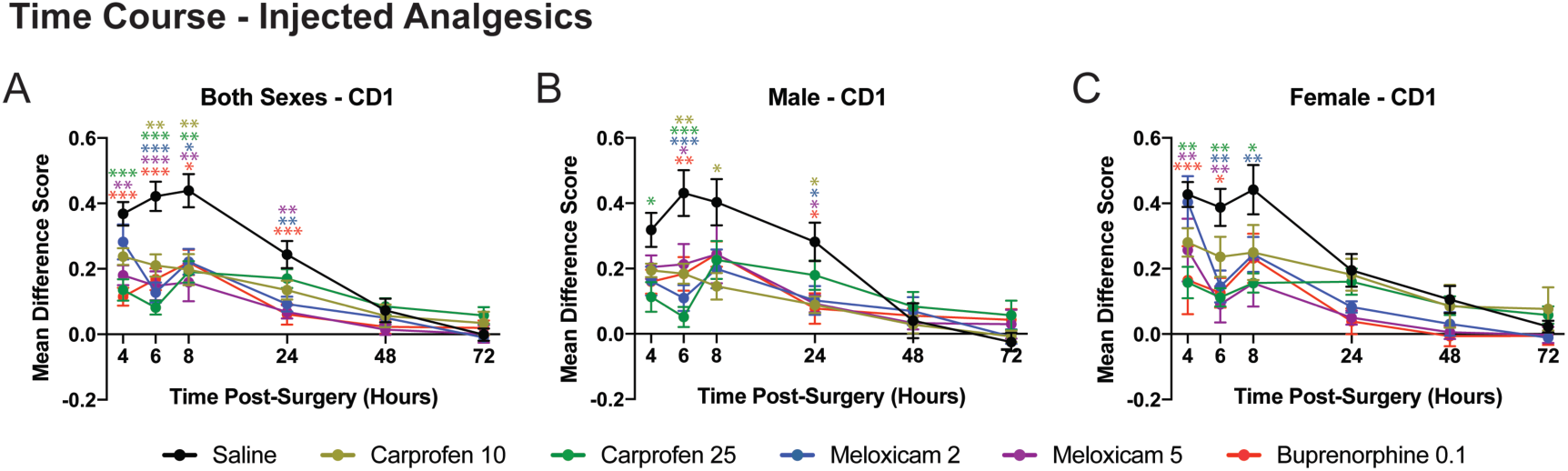
Time course postsurgery for injected drugs. Mean difference scores are shown for carprofen (10 and 25 mg/kg), meloxicam (2 and 5mg/kg) and buprenorphine (0.1 mg/kg) compared with saline (control) for pooled sexes (**A**), male mice (**B**), and female mice (**C**). Each drug has been color-coded as indicated by the legend and significance at each time point was determined by comparing with saline control. The color of the asterisks corresponds to the appropriate drug. * p<0.05; ** p<0.01, *** p<0.001.

### Analysis of water supply analgesics

Mean difference MGS scores for all mice in the water supply groups were calculated and plotted as shown in **Figure 3A**. A three-way ANOVA (sex x drug x repeated mea sures) revealed a main effect for drug (F_5,84_=2.82, p=0.02), while a main effect for sex (F_1,84_=3.55, p=0.06) and three-way interaction (F_25,84_=1.41, p=0.087) was approaching significance. To further examine the analgesic efficacy following craniotomy, separate one-way ANOVAs were performed at each time point. None of the drugs significantly reduced MGS scores at the 4 h (F_5,89_=1.32, p=0.26) and 6 h (F_5,89_=0.71, p=0.62) time points. However, buprenorphine reduced MGS scores at the 8 h time point (F_5,90_=2.36, p=0.046) and all drugs reduced MGS scores at the 24 h time point (F_5,90_=7.19, p<0.001) compared with control mice. No further differences were detected at 48 h (F_5,90_=1.18, p>0.05) or 72 h (F_5,90_=0.842, p>0.05). Overall sucralose consumption was not significant between drug conditions and likely not a contributing factor for differences in drug effects or lack thereof (one-way ANOVA, F_4,39_=1.42, p>0.05, **Figure 3A inset**). In addition, we estimated the average drug dose (in mg/kg) ingested by each mouse and these values were not significantly different from the injected doses when compared using multiple one-sample *t-tests* (**Table 2**). As was done for the injected analgesics, data were divided and plotted according to sex (**Figure 3B and 3C**).

**Figure 3.**
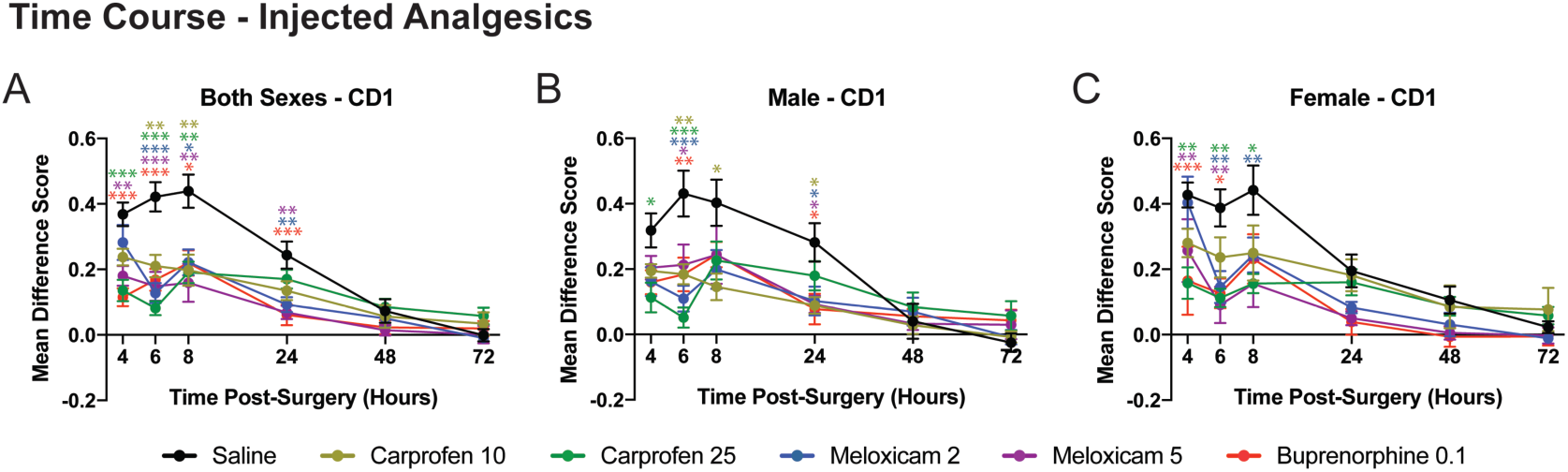
Time course postsurgery for drugs added to the water supply. Mean difference scores are shown for carprofen (10 and 25 mg/kg), meloxicam (2 and 5mg/kg) and buprenorphine (0.1 mg/kg) compared with saline (control) for pooled sexes (**A**), male mice (**B**), and female mice (**C**). Each drug has been color coded as indicated by the legend and significance at each time point was determined by comparing with saline control. The color of the asterisks corresponds to the appropriate drug. * p<0.05; ** p<0.01, *** p<0.001.

### Area Under Curve Analysis

To better compare the effect of drug treatment, sex and route of administration on postcraniotomy pain, we computed a single value (area under the curve; AUC) to represent each drug over the entire postoperative period. There was a main effect for drug (F_5,166_=12.44, p<0.001) and administration route (F_1,166_=13.88, p<0.001), but not sex (F_1,166_=0.037, p>0.05). However, the interaction between drug, administration route and sex was highly significant (F_5,166_=3.54, p<0.01). As shown in **Figure 4**, there were no major differences between drug treatment when administered through an injection or water supply for either sex. However, carprofen had a greater analgesic effect in female mice when given via the water supply compared with the injected route (10 mg/kg, p<0.001; 25 mg/kg, p=0.05).

**Figure 4.**
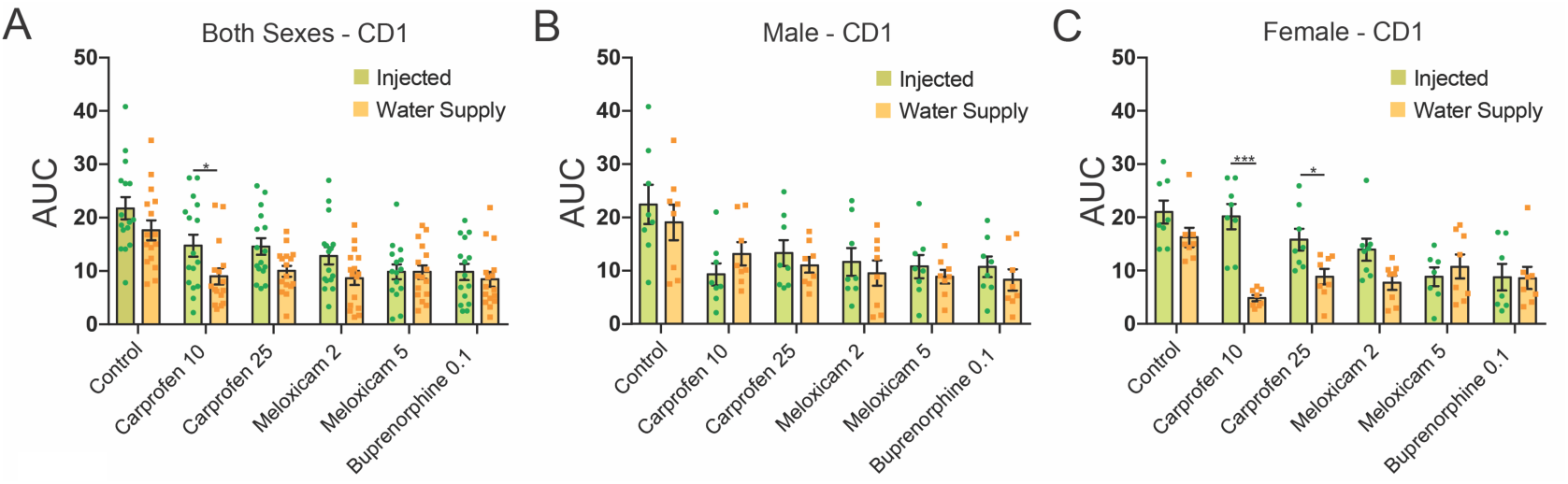
Comparison of AUC scores for control, carprofen (10 and 25 mg/kg), meloxicam (2 and 5mg/kg) and buprenorphine (0.1 mg/kg) over the entire 72 h postoperative period compared across administration route (injected vs. water supply). Pooled sexes (**A**), male mice (**B**) and female mice (**C**) are shown. Comparisons were conducted between administration routes for each panel, with lower AUC scores for female mice administered carprofen through the water supply. *p<0.05, **p<0.01.

In order to determine whether these effects generalized to another mouse strain, we assessed the AUC of MGS scores for C57BL/6N mice following craniotomy. We first noticed that the facial features of C57BL/6N were more difficult to distinguish than CD1 mice (**Figure 5**), however, orbital tightening, nose bulge, cheek bulge, and ear position were mostly visible. As such these action units were quantified and averaged to calculate the total MGS score for C57BL/6N mice. AUC was analyzed to remove the influence of time course and directly compare drug effects and administration route. Two-way ANOVA (drug x administration route) revealed a significant main effect for drug (F_5,82_=3.38, p=0.008), but neither administration route (F_1,82_=2.55, p=0.11) nor the interaction (F_5,82_=1.26, p=0.29) reached statistical significance. Carprofen (25 mg/kg) and meloxicam (5 mg/kg) significantly reduced the AUC when compared with control (**Figure 6A**; Tukey’s, p<0.05 and p<0.01, respectively), indicating an analgesic effect. However, the degree of analgesia by carprofen or meloxicam in C57BL/6N mice was less than CD-1 mice. This reduced analgesic effect of carprofen and meloxicam paralleled the overall lower MGS scores in the C57BL/6N control mice compared to CD1 mice (**Figure 6B**, t_45_=5.53, p<0.001).

**Figure 5.**
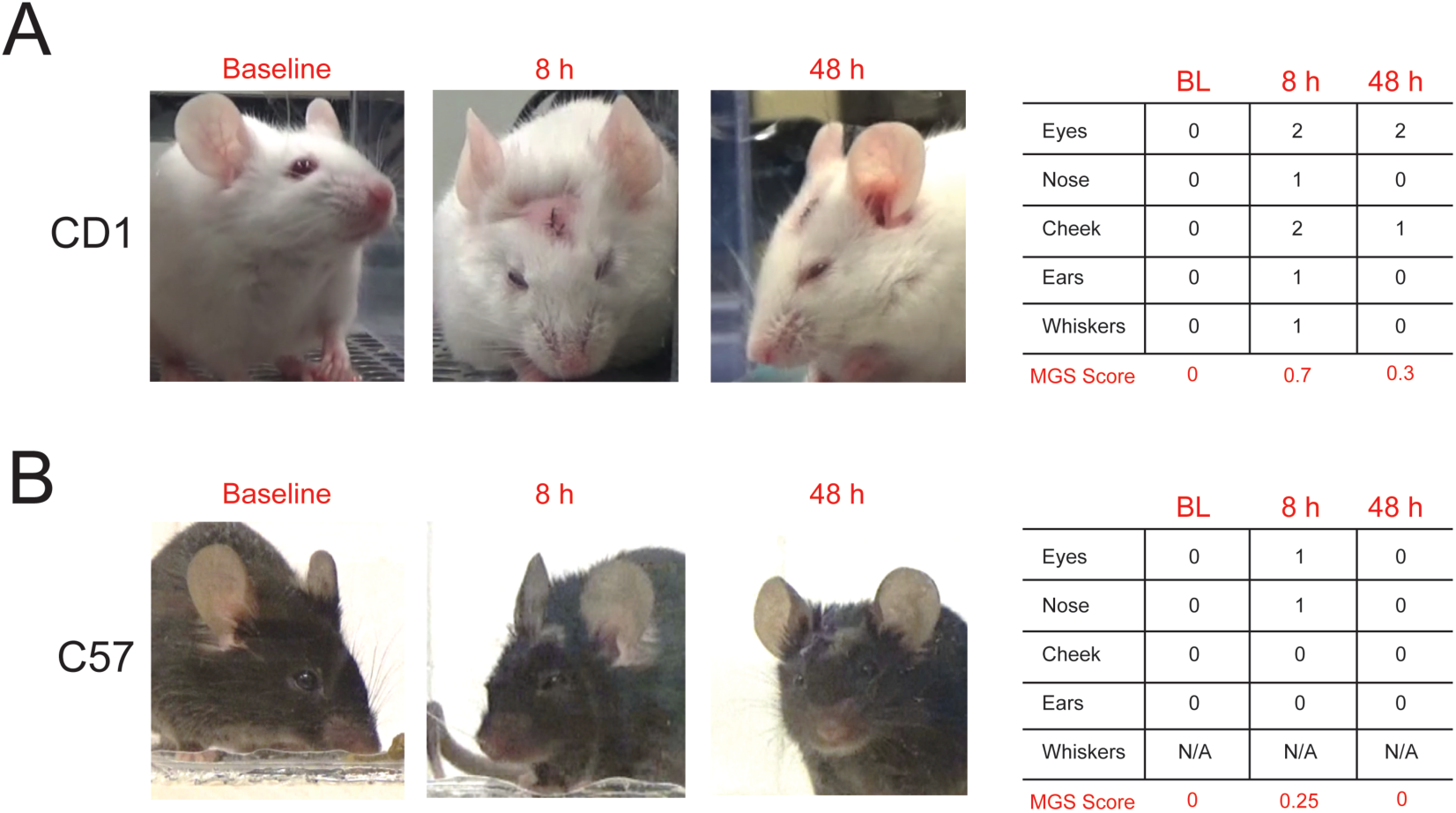
Representative images and corresponding scores for CD1 and C57BL/6N control mice at baseline, 8 h and 48 h postcraniotomy. CD1 images were scored using 5 action units, orbital tightening, nose bulge, cheek bulge, ear change and whisker change. All of the action units except for whisker change were scored for C57Bl/6N due to the poor detection of whiskers in the dark colored mice.

**Figure 6.**
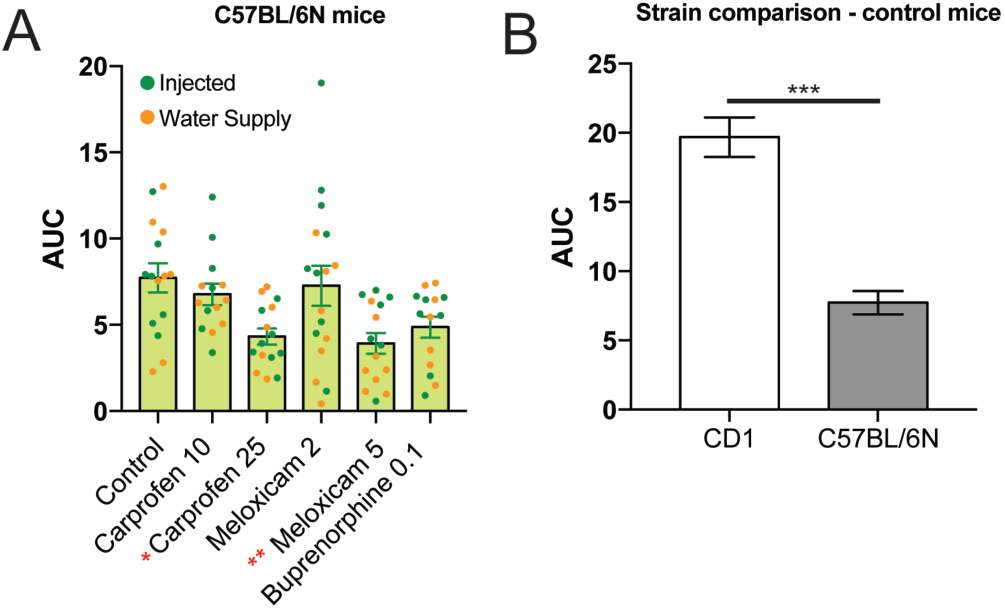
Area under the Curve (AUC) analysis for C57BL/6N mice. (**A**) AUC scores for C57BL/6N mice that were treated with carprofen (10 and 25 mg/kg), meloxicam (2 and 5mg/kg) and buprenorphine (0.1 mg/kg). For each condition, the scores of individual mice are presented as a dot plot with injectable analgesics indicated by the green circles and analgesics administered through the water supply as orange circles. In contrast with CD1 mice, MGS scores for the carprofen (25 mg/kg) and meloxicam (5 mg/kg) groups were significantly lower than control mice. (**B**) Overall higher MGS scores were found in CD1 control mice when compared with C57BL/6N mice. * p<0.05; ** p<0.01, *** p<0.001.

## Discussion

Unless postoperative pain is specifically being studied, pain control following surgery is an absolute requirement. The severity of pain following craniotomy has historically been viewed as minimal, but recent data suggests otherwise ^23^. This makes the detection of craniotomy-related pain in animals critically important for refinement, but since species and individual responses to pain are variable, evaluation and treatment of this pain are difficult. In the present study, we used the MGS to provide evidence that spontaneous pain following craniotomy lasted upwards of 48 h and commonly used analgesics reduced this pain when administered as an injected substance or through the water supply. We observed statistically significant reductions in MGS scores for all drugs given through both routes of administration, however, injected analgesics were more effective in the immediate hours (4 to 8 h) after surgery. Overall no sex differences were evident between drugs, but the NSAID carprofen was more efficacious for female mice when self-administered through the drinking water (**Figure 4**). In addition, we find that MGS scores were significantly lower for C57BL/6N mice following craniotomy, when compared with CD-1 mice and much higher doses of NSAIDS (carprofen or meloxicam) than currently recommended by our institution (**Table 1**) were required to reduce these scores.

**Table 1.**
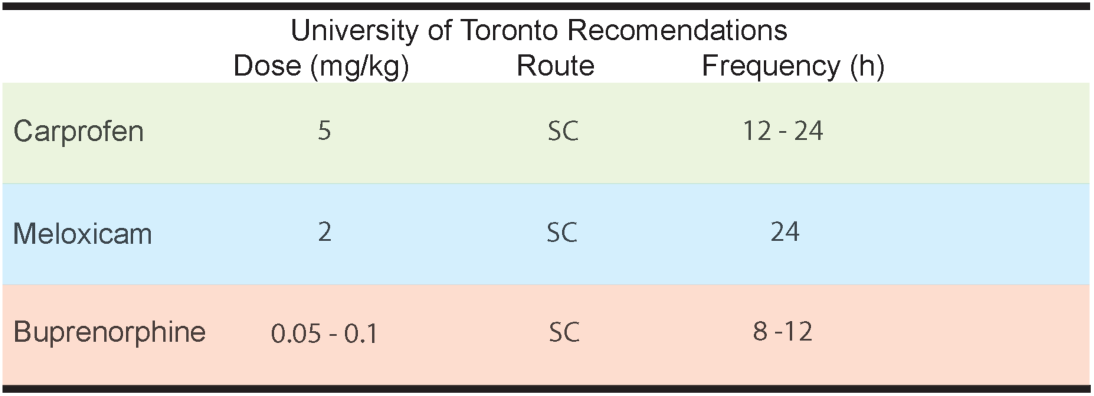
University of Toronto recommendations for dose, route and frequency of mouse analgesics following surgery.

In lab animals, the majority of analgesics are administered through an injectable formulation following surgery. However, recent trends and innovations have led to the development of alternative vehicles for the administration of drugs to reduce handling-related stress, such as gel formulations 24, sucralose in the water supply 25 or medicated pellets 26. We tested MediDrop(r) sucralose, a sweetened water gel for medication delivery (antibiotics, analgesics or experimental compounds), but did not find a clear advantage for analgesic administration following craniotomy when compared with equivalent injectable formulations – except when the NSAID carprofen was administered to female mice. Unfortunately, we did not measure serum concentration of drugs, but based on the consumption of water throughout the postoperative period, we estimate comparable levels of drug dosing for all mice independent of administration route (**Table 2**).

**Table 2.**
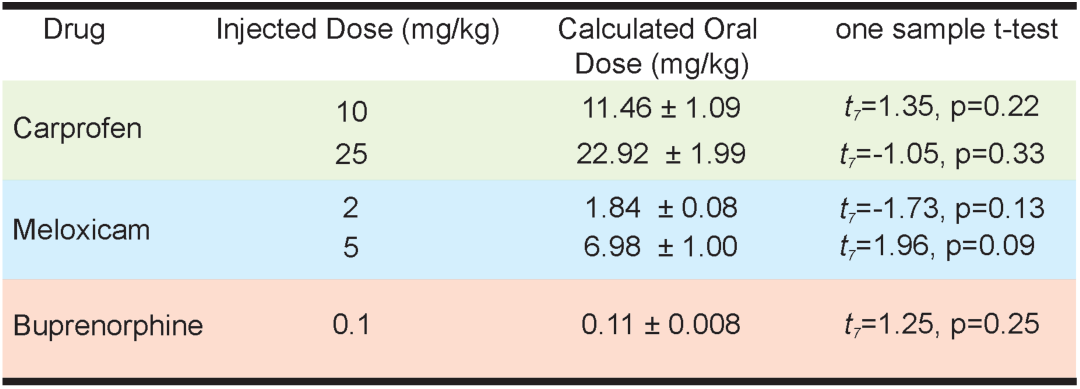
Estimated average drug dose (mg/kg) ingested compared to injected drug dose using a one-sample *t-test.*

Oral administration of analgesic drugs is intended to reduce the negative impact that restraint stress has on laboratory animals. Inherently, this is innovative because the stress associated with repeated physical restraint has been shown to negatively impact heart rate, body temperature, blood glucose and hormonal levels in mice see 27 for a review. In addition, restraint stress has been shown to delay cutaneous wound healing in mice 28, which may increase the risk of postsurgical infection and hemorrhage. The MediDrop(r) system – used here – may offer advantages, but its benefit may be small and details such as dose-response and consumption period prior to surgery may need to be refined. Since rodents have a natural aversion to new foods, sucralose was added to the water two days prior to surgery, this allowed mice to habituate to the novel taste and ensured that consumption remained constant throughout the postoperative testing period. We do not believe that prolonging the presurgical sucralose consumption period would have increased analgesic efficacy because previous studies have shown that 15 h of voluntary buprenorphine ingestion was required to reach similar or higher blood concentrations than subcutaneous administration 24. Previous studies have suggested that the willingness of the animal to ingest water may enhance inter-subject variation 29, but that does not apply here as there was comparable water consumption between all of our water supply groups (**Figure 3A inset**). Alternative approaches have been adopted to allow mice to voluntarily ingest analgesics, such as dissolving drugs in a sticky nut and chocolate paste (i.e. Nutella(r)), which mice readily consume, but this is not a sustainable solution as Nutella(r) has a high content of fat and sugar making it undesirable as a delivery vehicle for pain studies given the emerging literature on pain, inflammation and diet 30.

Finally, we believe that the current findings represent the most thorough assessment of postoperative pain and only investigation of craniotomy pain in rodents. The evaluation of different drugs, routes of administration, testing sites, mouse sex and mouse strain using the MGS as a primary output demonstrate the reliability of the MGS, but also point toward the CD1 mouse strain as an optimal choice for analgesic drug screening using this assay. The grimace scales have shown excellent consistency in pain detection among rodents and other species 11,12,14,17,21, however no single study has ever compared generalizability among experimenters, testing sites or pharmacological effects. Both the MGS and RGS (rat grimace scale) show robust inter-rater reliability, but these analyses have largely been conducted among individuals within the same test site and coding the same set of images. One consideration that we did not account for was the sex of the experimenter as experimenter sex has been previously shown to influence MGS scores 31, we did not find any test site differences even though the sex of the surgeon was different at each site. Given that husbandry practices may significantly alter mouse physiology and behavior, which may contribute to the lack of reproducibility in the literature, we did not make an effort to standardize practices among centers. Instead, we assessed the reliability and robustness of the MGS for craniotomy detection between centers and found no site-specific differences. This gives confidence in our results and supports the consistency of the MGS for detecting spontaneous pain following craniotomy despite differences in environmental conditions, experimental personnel and differences in equipment. Our results were less robust for craniotomy detection when the C57BL/6N mouse stain was used, but these mice had overall lower MGS scores than CD1 mice. Lower MGS scores may have produced a floor effect – especially where analgesics would be expected to decrease these scores – and it’s been previously reported that C57BL/6 mice have lower MGS scores than CD1 mice 15. Thus, our C57BL/6N results may be more reflective of the limited capability of MGS quantification in this mouse strain rather than lower overall pain in these mice. We believe that our findings have highlighted the need for postoperative analgesia following craniotomy, preferably in the form of an injected substance with buprenorphine remaining the most optimal overall choice.

## Methods and Materials

### Mice and test facilities

All experiments were approved by the appropriate animal care committees at the University of Toronto and conducted in accordance with the Canadian Council on Animal Care (CCAC) guidelines and the Ontario Animals for Research Act. In most experiments, naïve male and female CD-1 [Crl:CD1(ICR)] outbred mice (7 - 9 weeks of age) were tested in two different facilities. Mice tested at the University of Toronto Mississauga (*Martin lab*) were bred in-house or purchased from Charles River Laboratories. Mice bred in-house were housed with same-sex littermates in groups of 5 mice per cage at weaning. CD-1 or C57BL/6N mice tested at the University of Toronto St. George campus (Bonin Lab) were all purchased from Charles River. C57BL/6N mice (of both sexes) were assessed by the Bonin Lab to determine whether findings in CD-1 mice were generalizable to another commonly used mouse strain. Both facilities were temperature controlled (20 ± 1° C) with 12:12 h light:dark cycle and experiments were only conducted during the light period. Prior to the start of the experiments, all animals were separated and individually housed for a week. All mice had access to food (Harlan Teklad 8604) and water *ad libitum,* however, mice given drugs in water had their water supply switched to sucralose (MediDrop(r), Westbrook, ME) two days prior to surgery. Mice only underwent one surgery, and roughly equal numbers of male and female mice were tested in each cohort. For the C57BL/6N experiments, *n* = 4 mice per sex per group were used because no major sex differences emerged, and accurate detection of facial features was difficult in dark-colored mice, making any precise comparison between the two strains problematic.

### Digital video

Mice were placed individually on a tabletop in cubicles (9 × 5 × 5 cm high) with 2 walls of transparent acrylic glass and 2 side walls of removable stainless steel. Two high-resolution (1920 × 1080) digital video cameras (High-Definition Handycam Camcorder, model HDR-CX405, Sony, San Jose, CA) were placed immediately outside both acrylic glass walls to maximize the opportunity for clear facial shots. Video was taken for 30-min immediately before surgery (-1 day) and for 30-min periods centered around the postsurgical time points considered (4, 6, 8 12, 24, 48, and 72 h; see Figure 1 for an experimental timeline). The videos were subsequently analyzed, and one clear face shot per mouse was taken for every three minutes of video for subsequent analysis as previously reported 11. Average MGS scores for each mouse were calculated by averaging individual action unit scores across the entire duration of recording for each time point and mean difference scores were obtained by subtracting baseline MGS scores from the time point following surgery. For C57BL/6N mice, the brightness and contrast of all images was equally adjusted using Microsoft PowerPoint to better distinguish facial features. However, even with this image enhancement the coding of action units was difficult because of their dark fur color. Images were randomized across mice, time points and treatment, assigned random names, and scored in a blinded fashion.

### Surgery

A craniotomy, designed to mimic a neurosurgical procedure, was performed in isoflurane–oxygen-anesthetized mice by three surgeons (two in the Martin lab and one in the Bonin lab). Following anesthesia, mice were placed in a stereotaxic frame (BenchmarkTM, myNeuroLab) and administered a bolus of saline; mice in the injectable analgesic groups were also given their first treatment at this time. After shaving and disinfection of the surgical site, the skull was exposed following a 1.3-cm midline incision, and a 0.6 mm diameter circular craniotomy was performed over the skull region at coordinates AP = -2.46, ML = 0.55 from bregma. A needle (30G) was then used to remove the dura from the surface of the brain taking care to avoid puncturing the brain. The scalp was then sutured (^6-0^ vicryl, Ethicon) and mice were allowed to recover on a 37 °C heat pad with access to water and one pellet of food. Mice were monitored until ambulatory and then transferred to the experimental behavioral room for MGS assessment. Surgeons were blind to the animal treatment.

#### Injectable Administration of Drugs

Carprofen (10 and 25 mg/kg), meloxicam (2 and 5 mg/kg), and buprenorphine (0.1 mg/kg) were obtained from CDMV (St Hyacinthe, Quebec, Canada). They were dissolved in physiologic saline and administered at a volume of 10 mL/kg. Injections were given subcutaneously immediately before surgery and at 12 h time intervals following surgery. Doses were selected based on standard operating procedure and guidelines outlined by the University of Toronto, which considers craniotomy-related pain to be of mild-moderate severity. **Table 1**. outlines the University of Toronto dose recommendations for mouse analgesics following surgery, the preferred route of administration and frequency at which the analgesic should be administered. In most cases we selected doses at or above the upper range of these recommendations because previous studies have highlighted the inadequacy of recommended analgesic doses for treating postoperative pain in rodents ^14,16^.

### Oral Administration of Drugs

Carprofen (10 and 25 mg/kg), meloxicam (2 and 5 mg/kg), and buprenorphine (0.1 mg/kg) were dissolved in sucralose and made accessible to mice from 48 h prior to surgery. The amount of analgesic added to sucralose was calculated using the average sucralose consumptions of naïve CD-1 males (0.38ml/g body weight/day; n=8) and females (0.48ml/g body weight/day; n=8).

#### Statistical analyses

All statistical analyses were performed using SPSS version 20 (SPSS, Chicago, IL), with a criterion of α = 0.05. Normality and homoscedasticity of all data sets were confirmed by using the Shapiro–Wilk and Levene tests, respectively, and thus parametric statistics were used in all cases. Time-course data were analyzed using repeated-measures ANOVA, followed where appropriate by posthoc testing for repeated measures with Sidak correction for multiple comparisons. For initial analysis, time course data were separated according to the route of administration—that is, all drugs were first compared based on administration route (i.e. injectable or water supply). In addition, data were further quantified with respect to the area under the time course curve (using the trapezoidal method) over the postoperative testing period (72 h in all cases). The area under the curve data were analyzed using either a one-, two-, or three-way ANOVA followed by Dunnett’s case-comparison posthoc test. Calculated drug doses for the oral administration groups were analyzed by one-sample Student *t* test, comparing the calculated with the actual injected dose of each drug. The routes of administration were directly compared for all drugs through area under the curve analysis.

## Acknowledgments

We thank Jasmine Hg, Annissa Ho, and Jerry Li for their assistance with the data organization and coding. This research was supported by the Canada Research Chairs Program (L.J.M. and R.P.B.), American College of Laboratory Animal Medicine (R.P.B, the University of Toronto Centre for the Study of Pain (R.P.B), the Research Scholars and Activity Fund, University of Toronto Mississauga (L.J.M.) and the Natural Sciences and Engineering Research Council of Canada (I.L.). Chulmin Cho was partially supported by a trainee fellowship from the University of Toronto Centre for the Study of Pain

## Author Contributions

L.J.M and R.P.B conceived of the project. C.Cho., V.M., I.L., C.C., M.L., M.D., C.P., and P.U. collected data. C.C., D.H., and L.J.M. analyzed data.

L.J.M. wrote the paper with input from C.C. and V.M. All authors edited and commented on the paper.

